# Three-dimensional spatial transcriptomics uncovers cell type dynamics in the rheumatoid arthritis synovium

**DOI:** 10.1101/2020.12.10.420463

**Authors:** Sanja Vickovic, Denis Schapiro, Konstantin Carlberg, Britta Lötstedt, Ludvig Larsson, Marina Korotkova, Aase H Hensvold, Anca I Catrina, Peter K Sorger, Vivianne Malmström, Aviv Regev, Patrik L Ståhl

## Abstract

The inflamed rheumatic joint is a highly heterogeneous and complex tissue with dynamic recruitment and expansion of multiple cell types that interact in multifaceted ways within a localized area. Rheumatoid arthritis synovium has primarily been studied either by immunostaining or by molecular profiling after tissue homogenization. Here, we use Spatial Transcriptomics to study local cellular interactions at the site of chronic synovial inflammation. We report comprehensive spatial RNA-seq data coupled to quantitative and cell type-specific chemokine-driven dynamics at and around organized structures of infiltrating leukocyte cells in the synovium.

---

Rheumatoid arthritis (RA) is a chronic autoimmune disease that primarily affects the joints. It consists of two broad subsets, seropositive and seronegative. Seropositive RA, comprises two thirds of patients, who often exhibit more severe symptoms, is a classical autoimmune disease defined by the presence of rheumatoid factor (RF) or anti-citrullinated protein antibodies (ACPA)^1^. RA pathogenesis involves complex interactions between fibroblasts and cells of the innate and adaptive immune systems that lead to imbalanced secretion of pro- and anti-inflammatory cytokines^2^. Studies of RA pathology have reported markers for an activated synovial fibroblast state^3,4^, while others revealed the contribution of adaptive immune responses in isolated MHC class II-dependent T cells in response to the production of specific cytokines^5,6,7^. Activation and expansion of fibroblasts in the synovial lining also contributes to changes in the extracellular matrix, further contributing to bone and cartilage destruction^8^. Current existing therapies, mainly targeting the immune cell components, can reduce symptoms and progression, but only ~60% of patients respond adequately to these treatments^9^.

Regions within sites of inflammation are filled with local accumulations of infiltrating leukocytes that form more or less organized structures. Such aggregates histologically resemble secondary lymphoid organs (SLOs) and are often termed tertiary lymphoid organs (TLOs)^10^. Patients with large and developed TLOs have been reported to respond more poorly to existing therapies^11^, but this is a topic of discussion in the field^12,13^. Recently, single cell RNA-Seq studies have uncovered additional fibroblast and immune cell types and states associated with RA and TLOs^14,15^. However, the spatial organization of these cells and their impact on TLO pathogenesis in RA remains unknown.

We have previously developed Spatial Transcriptomics^16–18^ (ST), a method for high-throughput transcriptome profiling that retains spatial information in tissues^16^. In ST, transcriptomes are barcoded directly from frozen tissue sections. Tissue sections are placed on a glass slide covered with 1,000-2,000 features, each carrying a uniquely barcoded poly(d)T capture sequence enabling spatial mRNA capture. Tissue sections are then stained with Hematoxylin and Eosin (H&E) and imaged by transmitted light microscopy, followed by gentle permeabilization, mRNA capture on the poly(d)T probes and RNA-seq. Analysis of the resulting data provides a direct link between histology and RNA-seq.

Here, we used ST to spatially profile synovial tissues from seropositive and seronegative RA patients. To address the genomic variability and profile the TLO-like structures, we have studied gene expression as localized (2D) and three dimensional (3D) views. We report the resulting gene expression signatures, quantitative single-cell morphological features and patterns of cell migration patterns at the sites of synovial inflammation. This provides the first 3D, high-throughput transcriptomic view of rheumatoid arthritis-affected synovial biopsies.

## Results

### 3D spatial profiling of RA synovia

To study the spatial organization in RA, we profiled 23 tissue sections by ST from five biopsies collected from RA patients at the time of joint replacement; specimens comprise three knee and two hip biopsies (**Fig. 1, Supplementary Table 1**). We optimized the technology for the tissue with the specific characteristics of synovia (**Methods, Supplementary Fig. 1**), collected profiles from consecutive sections, and aligned and interpolated the data to create a 3D view within each biopsy (**Methods, Fig. 1**). This 3D sampling approach spanned larger distances creating the first exploratory multidimensional view of an RA synovial tissue biopsy.

**Fig. 1.**
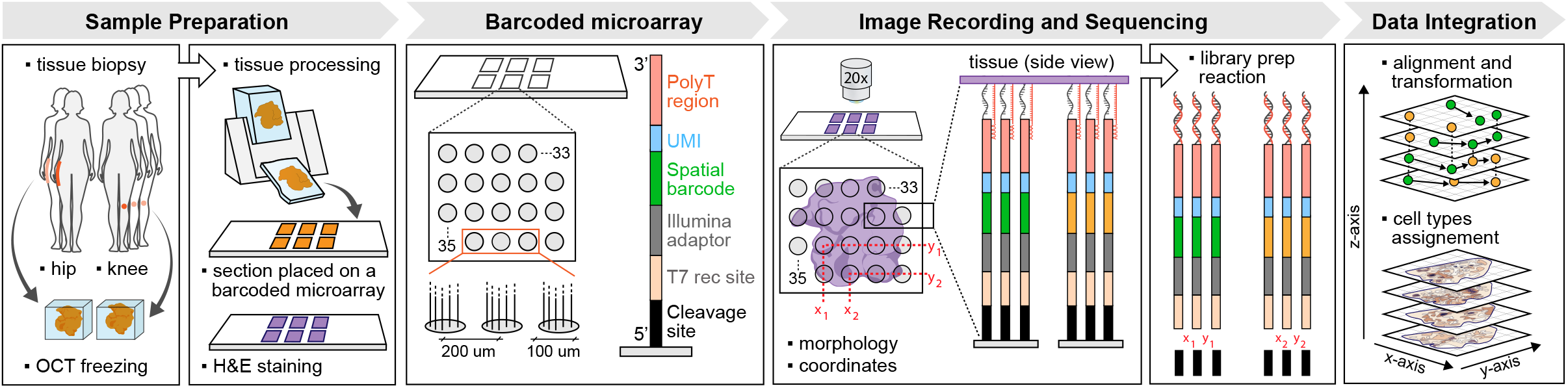
Sampling and spatial barcoding of rheumatoid arthritis samples. Synovial tissue from two sites, hip and knee, was sampled and the biopsies cryopreserved in OCT compound. The biopsies were cryosectioned and placed on a spatially barcoded microarray. Tissue sections were H&E stained and the images recorded. While recording histology, positional information of each spatial (x,y) feature was also tracked. Cells in the tissue were gently permeabilized and mRNA molecules captured on the spatially barcoded poly(d)T capture probes. The cDNA synthesis reaction was performed on the slide surface and mRNA information copied. Libraries were prepared and pair-end sequenced. The data was processed so that spatially barcoded expression information and the morphological images were registered and aligned. This resulted in spatial data transformation, interpolation and imminent visualization.

### Variation within and between TLO-like structures in RA

The biopsies from RA joints contained regions where infiltrating leukocytes (“infiltrates”) organize into cell-dense areas to form TLO-like aggregates^19^ (**Fig. 2a, left**), which we annotated manually. We also detected TLOs automatically as regions of high density and distinct cellular topology (**Methods, Supplementary Fig. 2**). 80% of all manually-annotated infiltrates were in regions with a cell density score higher than 70% (**Supplementary Fig. 3**).

**Fig. 2.**
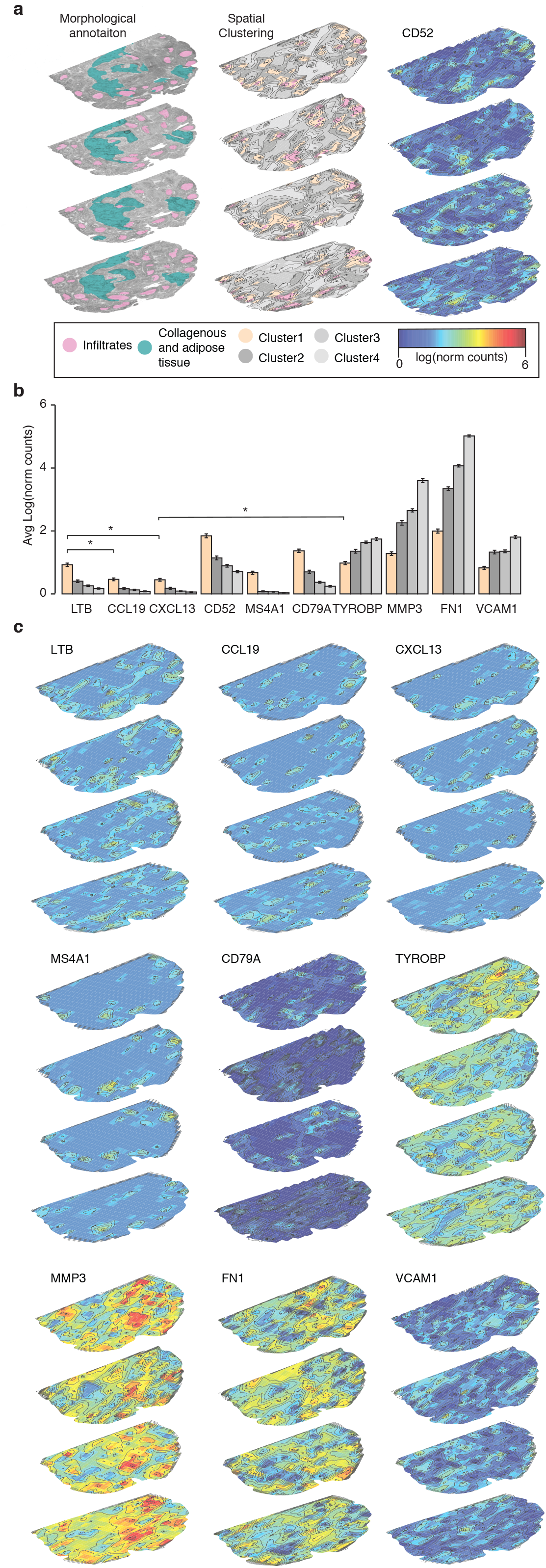
Spatial data clustering in RA1. (**a**) Morphological annotation, spatial clustering (color code) and CD52 spatial expression (color scale). Pink marks spatial infiltrate positions that overlap between the morphological annotation and spatial clustering (Cluster1). (**b**) Average expression of genes found in different spatial clusters. Statistical significance markings (Wilcoxon’s rank sum test) are displayed; 0.005<p≤0.05 (*). (**c**) Spatial expression of nine differentially expressed genes as determined by clustering. Color-scale denotes gene expression and is shared between panels (a) and (c). Color code is shared between panels (a) and (b).

We then looked for differences between and within infiltrates in one biopsy (RA1, RF^+^ACPA^+^ patient, knee, **Methods**). Analysis of spatially variable gene expression patterns revealed two clusters of infiltrate features in the TLO-like aggregates (**Methods, Supplementary Fig. 4, Supplementary Table 2**) varying in the expression of multiple genes including CD52, MS4A1 and FN1 (**Supplementary Fig. 5**). These differences were found not only between aggregates but also within aggregates along its z-axis, which we could capture due to 3D sampling.

### Cytokine signaling from spatially resolved profiles of the RA synovium

Next, using unsupervised clustering of the regions in the entire RA1 biopsy, we identified four major spatial domains: Cluster 1 included 87% of all annotated RA1 infiltrate data points (**Fig. 2a, middle, Supplementary Table 3**), and Clusters 2-4 included the remainder and followed radial spatial patterns at consecutively increasing distance from the infiltrate Cluster 1 regions and had lower cell density scores (**Fig. 2a**).

In the RA1 Cluster 1 infiltrates, lymph node/TLO-associated genes (LTB and CCL19) and genes associated with B cells, T cells and their cross talk (CXCL13, CD52, MS4A1 and CD79A), were up-regulated (Wilcoxon’s rank sum test, p≤0.05), both as averaged signatures per spatial cluster (**Fig. 2b**) and as high-resolution spatial maps (**Fig. 2c**). CXCL13/CCL21 expression has previously been associated with formation of the spatial niches of T cells in model systems^20^ and CXCL13 is also a key chemokine produced by T follicular and T peripheral helper (Tfh and Tph) cell subsets used in promoting B-cell mediated responses^21^. CXCL12/CCL19 expression, on the other hand, affects the spatial distributions of dendritic (DCs), B and plasma cells in TLOs^20^. Signaling driven by these cytokines has also been previously associated with overexpression of LTA and LTB^20^, a finding recapitulated in our spatially resolved data (Wilcoxon’s rank sum test, p≤0.05, **Fig. 2b-c**). In addition, we find downregulation of CXCL13 in TYROBP-high areas (present in 46% of all spatial Cluster 1 features) (**Fig. 2b-c, Supplementary Fig. 4b**). TYROBP-mediated ITAM pathway activity has previously been associated with immune cell co-modulation of bone cells in RA^22^.

In areas neighboring TLO-like aggregates (Clusters 2-4), gene expression was characterized by significantly increased (Wilcoxon’s rank sum test, p≤0.05) levels of metalloproteinases (MMP3, **Fig. 2c**) which are involved in extracellular matrix degradation. Fibronectin-1 expression (FN1, **Fig. 2c**), an extracellular matrix protein expressed by fibroblasts that induces transforming growth factor-beta secretion, and vascular cell adhesion molecule 1 expression (VCAM1, **Fig. 2c**), were also upregulated, supporting the hypothesis that newly recruited hematopoietic cells are retained in the TLO-like structures^23^.

The spatial organization generalized by 3D ST profiling of consecutive sections in another joint-affected RA patient biopsy (RA2, RF^+^ACPA^+^ patient, hip), with large infiltrates that spanned most of the sampled area (**Supplementary Fig. 6a**). Unsupervised clustering partitioned the regions to three major clusters having distinct spatial expression patterns (**Supplementary Table 3**). Cluster 1 corresponded to the infiltrate areas (**Supplementary Fig. 6a)**, comprising 90% of regions annotated by cellular morphology and high cell density (**Supplementary Fig. 7**), and the two other clusters formed a radial pattern. Key genes followed similar patterns to those in the RA1 sample, and included induction of CD52, LTB, CCL19 and MS4A1 infiltrates (Cluster 1, Wilcoxon’s rank sum test, p≤0.05) and increased FN1, MMP3 and PRG4 expression in the surrounding areas (Clusters 2-3, Wilcoxon’s rank sum test, p≤0.05, **Supplementary Fig. 6b-d**).

### Dense volumetric analysis highlights chemokine driven T and B cell organization in TLO-like aggregates

Closer examination of intra-infiltrate spatial patterns, distinguished T and B cell specific variation within the infiltrates (**Fig. 3, Supplementary Table 2**). For example, following infiltrate 6 (**Fig. 3, Supplementary Fig. 6a**) in 3D, we observed co-expression of CD52 and MS4A1 in highly localized patterns within TLO-like aggregates. Upregulation of CCL21/CCL19 in RA2 (present in 75% of all Cluster 1 features), was accompanied with high expression of IL7R in 39% of spatial measurements (**Fig. 3**). In combination with CXCL13 upregulation in RA1, these data suggests a process of self-organization of T and B cells in TLO-like aggregates. We also reproduced our observation that CXCL13 is downregulated in TYROBP-high areas, which were present in 53% of all TLO-like spatial features. The CCL21^+^ sites were restricted to the centers, while MZB1^+^XBP1^+^ sites were spatially overlaid with the outer rim of the TLO-like structures (**Fig. 3**), suggesting localized prevalence of plasma cells on the TLO-like edge sites. Similar analyses of the three additional samples (RA3-5) *i.e*. spatial and intra-infiltrate clustering was performed (**Supplementary Fig. 8a, Supplementary Tables 1-3**). RA3, a specimen from a patient having clinical characteristics similar to those of RA1-2 (RF^+^ACPA^+^), also exhibited significantly higher (Wilcoxon’s rank sum test, p≤0.05) expression of CD52 and MS4A1 in Cluster 1 that marked the cell dense infiltrate regions. These marker genes were also significantly higher (Wilcoxon’s rank sum test, p≤0.05) in the same cluster denoting the TLO-like structures in RA4-5 (RFACPA^-,^, **Supplementary Fig. 8b**) although the overall expression of these markers was significantly lower (one-sided *t*-test, p≤0.05) in the RF^+^ACPA^+^ (seronegative) than in RF^+^ACPA^+^ (seropositive) patients. In regions surrounding Cluster 1, RA3 was again similar to the other two seropositive patients with upregulation of MMP3, FN1, PRG4 and TYROBP in Clusters 2-3 (Wilcoxon’s rank sum test, p≤0.05). In seronegative patients, we did not detect the same gene expression patterns (**Supplementary Fig. 8b**). There, the same genes were instead found to be downregulated in the areas surrounding the TLO-structures (Wilcoxon’s rank sum test, p≤0.05). CCL19, which we and others reported as implicated in B cell distribution in the TLO sites, was also found to be significantly downregulated (Wilcoxon’s rank sum test, p≤0.05) in the TLO-structures of seronegative patients. This was also the opposite of what was observed in the three seropositive patients (**Supplementary Fig. 8b**).

**Fig. 3.**
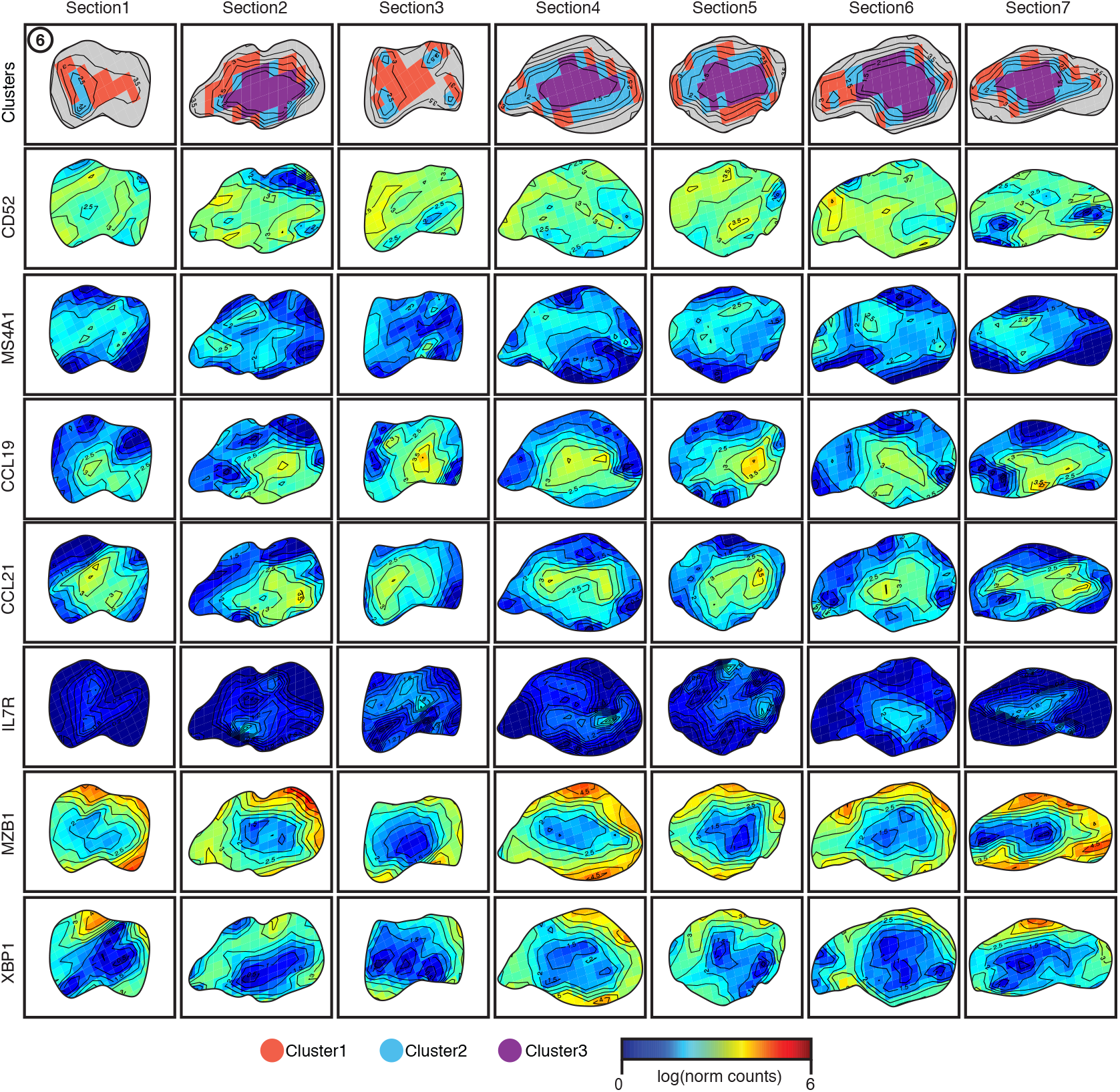
RA2 infiltrate dynamics. Zoomed in expression (color scale) of spatial clusters followed by seven example genes (rows) in the Infiltrate6 region across in RA2 sections (columns).

### Substantial variation in cell composition and spatial organization in the RA synovium

To relate the spatial profiles to the cellular composition of RA regions, we used a previously published scRNA-seq reference^14^ to define cell type specific signatures, and scored our spatial regions in each of the five patients (**Fig. 4, Methods**). Out of the 13 cell types available in the reference^14^, plasma cells, macrophages, CD55^+^ fibroblasts (F1) and THY1^+^ fibroblasts (F2B) were found in every analyzed sample and section (**Supplementary Fig. 9**), while B cell abundances were significantly higher in the tissue volume of RA2 than in any other sample (one sided t-test, p<0.05), with similar significantly spatially enriched (one sided t-test, p<0.05) Tph cell type distributions in RA1 and DCs distributions in RA4.

**Fig. 4.**
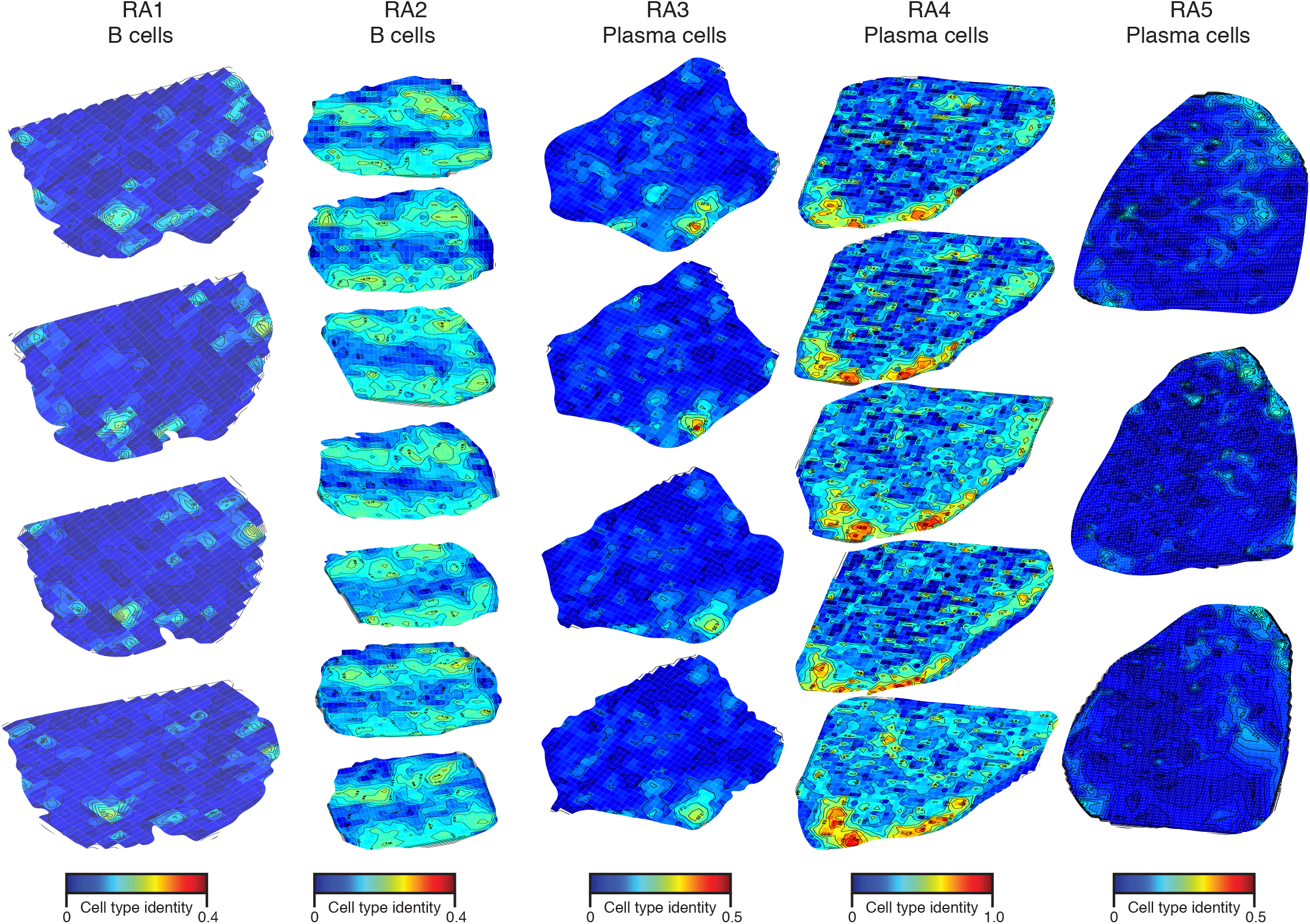
Spatial distribution of cell types in the rheumatoid arthritis synovium. Most abundant cell types (color scale) shown in each of the five RA patient samples (columns) and across all spatially prof **Supplementary information for:**

In RA1-3, macrophage-enriched cell areas were on average significantly co-localized with higher presence of F2B fibroblast cells in the whole tissue volume (Pearson’s R 0.93; 0.80; 0.72, p<0.05, respectively RA1-3, **Supplementary Fig. 10a**). Additionally, in specific structures spanning both TLOs and surrounding areas in RA1-2, macrophage areas were found together with plasma cell areas (**Supplementary Fig. 10b-c**). In RA3, we observed a trend in which macrophage-rich areas were spatially correlated with F2B-rich areas in 2 out of the 9 TLO-like structures (Pearson’s R 0.99, p<0.05, **Supplementary Fig. 10d**), whereas F2B-rich areas spatially surrounding the TLO-structures were exhausted of plasma cells (data not shown, Pearson’s R −0.98, p<0.05). Interestingly, while the TLO-like structures in RA1 and RA2 were dominated by both B cells and CD4^+^ T cells, RA2 was again specific with significantly higher (one-sided *t*-test, p<0.05) abundances of CD8^+^ T cells and Tph cells. In RA3, CD8^+^ T cells and Tph cells were not detected in the tissue and only few B cells were detected (**Supplementary Fig. 9**). No significant levels (one-sided *t*-test, p>0.05) of either B or T cell scores were seen in RA4-5 in the tissue volume. Conversely, DCs were substantially increased in RA4-5 and not contained to specific areas in the tissue volume nor was their expression spatially correlated to B cell presence. This is the opposite of what was observed in RA1, in which tissues recruitment of DCs in areas surrounding the infiltrates was associated with a decrease in B cells (Pearson’r R - 0.68, p<0.05, **Supplementary Fig. 10e**). In RA2, which had the largest TLO-like structures and most B cells, DCs were few, their abundance significantly lower (one-sided t-test, p<0.05) as compared to all other tissue volumes but these DCs were also spatially contained to B cell sparse zones (Pearson’r R −0.60, p<0.05, **Supplementary Fig. 10e**).

### Connecting H&E and spatial transcriptomics reveals unified spatial clusters of morphological and molecular profiles features

Connecting morphological data to tissue-specific molecular profiles^24–27^ helps translating clinically relevant H&E information^28,29^ to spatially resolved molecular signatures. We hypothesized that distinct cellular morphological features would also be reflected in different ST profiles and in other spatial features, such as cell density. To explore this, after cell segmentation of the H&E image accompanying spatial transcriptomics, we clustered the segmented cells by their morphological features (**Methods, Supplementary Fig. 11a**) and then examined their relation to other features from the H&E image and from ST.

Cluster1 (RA2), which represents the areas of infiltration, was enriched in specific H&E-derived cell clusters (**Supplementary Fig. 11b**), and those were, as expected, also regions of high cellular density of small cells across all samples (**Supplementary Fig. 11c,d**). Conversely, Cluster4 (in RA2) was prevalent in other H&E-defined cell clusters and those were associated with phenotypically large cell sizes (**Supplementary Fig. 11d**), in line with abundances of larger cell types like macrophages and fibroblasts in those distinct areas (**Supplementary Fig. 11e-f**). Across all sections, we distinguished quantitative descriptions of cellular morphology and architecture in TLO-like areas and related them to single cell signatures (**viewable in** an extension we developed for histoCAT^24^, **Methods**).

## Discussion

Spatially resolved genomic analysis of disease tissue holds promise for better precision phenotyping of patients and assessment of treatment responses in a manner that combines established histopathology with comprehensive molecular profiling. Here, we created an exploratory 3D spatial gene expression catalogue comprising high-resolution transcriptome-wide volumetric maps correlated to morphological features. This serves, to the best of our knowledge, as the first combined morphological, spatial and transcriptional blueprint of tissue from autoimmune disease patients, and spans multiple sections from five patient specimens.

The spatial clusters observed in synovial biopsies were distributed radially around the infiltrate sub-regions, further confirming the uniqueness of signals and cell types present in those areas, and highlighting the potential role of complex center-based TLO-like dynamics in these biopsies. The spatial cell type organization throughout the 3D volume was transcriptionally connected to genes related to extracellular structure organization, regulation of B cell activation and proliferation, cytokine production and platelet degranulation in all analyzed samples.

Tph cells initiate B cell to plasma cell differentiation^30,31^ and given CXCL13 and IL-21 production, recruit more B cells to the TLO sites resulting in increased localized autoantibody and cytokine production^32^. Interestingly, 34% of these regions expressing Tph signature genes included RASGRP2 overexpression in the RA2 seropositive patient tissue, a gene previously reported in the development of destructive arthritis^33^. Fibroblast cells surrounding TLOs have, on the other hand, been associated with propagation of T cell-rich zones and are considered marker features of lymphoid neogenesis^34^. The CD55^+^ fibroblast population was present in the synovial lining (*i.e*. outer rim of the tissue) while CD90^+^ (F2B) fibroblast populations were located closer to the TLO regions in all seropositive samples. Seronegative tissue volumes lacked robust signals of ongoing adaptive immune responses and were characterized by increased presence of DCs. DCs are involved in recruiting proinflammatory immune cells including macrophages, neutrophils and monocytes in RA^35^. Specifically in the seronegative tissue volumes, we report similar spatial cell distributions - the fibroblast populations as well as macrophages were significantly overexpressed (one-sided t-test, p<0.05) in the TLO structures; implicating a completely different immunological ‘drive’ in the sites of inflammation as compared to spatially deconvolved disease responses in seropositive tissues.

Combining morphological features and high-throughput spatial signatures could aid in clinical diagnosis and overall disease management of RA. ST technology is compatible with conventional histological staining, has fast turnaround times and user-friendly laboratory setup. Future clinical studies using high-throughput spatially resolved transcriptomics^36^ may be able to provide higher statistical power and more insights into monitoring disease severity and treatment-specific responses in seropositive and seronegative rheumatoid arthritis.

## Methods

### Patient information and sample collection

Synovial tissue biopsies from knee or hip joints were obtained during orthopedic replacement surgery. Additional patient information can be found in **Supplementary Table 1**. Ethical approvals were granted by the Ethics Committee of Karolinska University Hospital (2009/1262-31/3) and patients gave their informed written consent to participate in the study. The biopsies were snap frozen in isopentane prechilled with liquid nitrogen within 15 minutes of collection and kept at −80°C until embedding in OCT (Sakura, The Netherlands) and sectioning could be performed.

### Spatial transcriptomics

Tissues were cryosectioned at 7μm thickness. Each section was carefully handled inside a cryotome (CryoStar NX70, Thermo Fisher Scientific, Life Technologies, Paisley, UK) and placed onto an individual array without any direct contact between the array surface and the cryotome as to avoid contamination. All sections were placed in the same fashion onto individual arrays. RA1 sections were sectioned at 21μm distance from each other while the RA2-5 sections were consecutives (z=7μm). The whole slide was then warmed for 1 min at 37°C and immediately fixed for 10 min at room temperature (RT) in a 2% formaldehyde solution (1:20 37% formaldehyde acquired from Sigma-Aldrich, Missouri, USA in 1x PBS pH 7.4). The sections were dried with isopropanol and stained with hematoxylin and eosin (H&E). To ensure proper staining, the dried sections were incubated for 7 min with hematoxylin (Mayer’s solution, Sigma-Aldrich, Missouri, USA) followed by 2 min in bluing buffer (DAKO, Agilent, California, USA) and 10 sec in eosin Y (1:20 in slightly acidic pH 6 Tris). To record both morphological and positional information, each tissue area was imaged at 20x resolution (Olympus, Japan) individually with a Metafer system (MetaSystems, Germany). Image stitching was performed using VSide software provided by MetaSystems. The Spatial Transcriptomics protocol was carried out as previously described^17,37,38^. Sequencing was carried out on either the Nextseq 550 (RA1-2) or Novaseq 6000 (RA3-5) instruments.

### Data mapping, annotation and filtering

Data was pre-processed using a recently published pipeline^39^. Raw sequencing reads were demultiplexed using CASAVA according to the TruSeq LT index information. The forward read contained 28-30 nt; 18 nt spatial barcode followed by a semi-randomized 9 nt unique molecular identifier (UMI) (RA1-2) or randomized UMI (7 nts, RA3-5), while the reverse read contained the 50 nt transcript information. The first five bases in the reverse read were hard trimmed and then the rest of the read was quality trimmed based on the Burrows-Wheeler aligner. Trimmed reads were mapped to the human genome reference (GRCh38) using STAR^40^. Mapped reads were annotated based on Ensembl’s v79 information and then paired with their forward read, UMI-filtered with a Hamming distance of 2 and counted using HTseq-count^41^. Quality control statistics were computed as number of paired reads per spatial barcode; number of UMI counts per spatial barcode and number of unique gene counts per spatial barcode. Data were normalized per biopsy using a linear regression approach^42^ with a mean gene cutoff per ST spot prior to normalization as recommended by the developer.

### Image registration, alignment and visualization

Images were randomly down-sampled to approximately the same image size per patient biopsy. In the RA1 biopsy, all sections were also cropped to contain approximately the same tissue areas. Image background was removed using scikit-image^43^ before registering the sections using SCALED_ROTATION (biopsies RA1 and RA2) and RIGID_BODY (biopsies RA3, RA4 and RA5) from PyStackReg^44^. Transformation matrices were used to align the spatially resolved count matrices in the same fashion. All of the following data processing was performed in R. As the spatially resolved data is of restricted resolution, the data was interpolated using the akima package in R over the tissue section area to aid in data interpretation. Volumetric expression heatmaps were created that could be viewed interactively using the developed RShiny application (https://spatialtranscriptomics3d.shinyapps.io/3DSeTH/).

### Single cell segmentation

Single cell segmentation was performed by combining Ilastik 1.3.2^45^ and CellProfiler 3.1.8^46^. Random forest classification implemented in Ilastik was used to train three distinct classes (nuclei, membrane, and background) to enable the prediction and export of probability maps. CellProfiler was then used to segment those exported probability maps to create labeled single cell masks for downstream analysis.

### Coupling single cell topology to ST data

ST 100μm barcoded area locations were used to crop areas of 200×200 pixels from the corresponding H&E images. These cropped and segmented images and imported into histoCAT^24^ for single cell quantification and spatial analysis. ST based phenotypic clusters were matched to the single cell data as well as the manual infiltrate annotations. Each image was saved as an individual interactive session for histoCAT loading.

### Phenotyping cell type calling

We used PhenoGraph^47^ with the code in https://github.com/jacoblevine/PhenoGraph to define phenotypic groups (PG) based on the morphological single cell readouts. We used histoCAT^24^ to extract mean marker expression as well as morphological features from the single cell mask. The default setting (30 nearest neighbors) was used to define 25 distinct phenotypic groups using a fixed seed for the Louvain method (random seed: 2).

### Spatially resolved DE analysis

To cluster regions, most variable genes were selected as previously described^42^ and principal component analysis (PCA) performed on the sub sampled and normalized *region x gene* expression matrices, followed by two dimensional *t*-stochastic neighbor embedding (tSNE)^48^. Hierarchical clustering was done on the 3D tSNE reduced data to determine numbers of individual clusters present in the whole tissue volume followed by differential expression (DE) analysis using a likelihood ratio test^49^. DE genes between the clusters were called as differentially expressed^18^ if satisfying the following criteria: p<0.001 and log ratio >0.5.

### Single cell signatures

Single cell type signatures were downloaded from Stephenson *et al^14^*, and the top 200 markers *m_l_* were kept for each cell type *l* with the following criteria: average log fold change>1 and FDR<5%. A total of 13 cell types were present in the reference. ST matrix is defined as *region x gene* matrix for a total of *i* regions and *j* genes. To score each cell type *c_l, r_* assignment per each individual spatial feature *S_i,j_*, the normalized ST matrix for was first subset for *m_l_* if more than 3 *m_l_* genes for each *S_i,j_* were present; creating a *R x K* matrix. Then, we computed the correlation coefficient over each *S_i,j_* for each pair of genes (*j, k*) and a total of R regions such that:

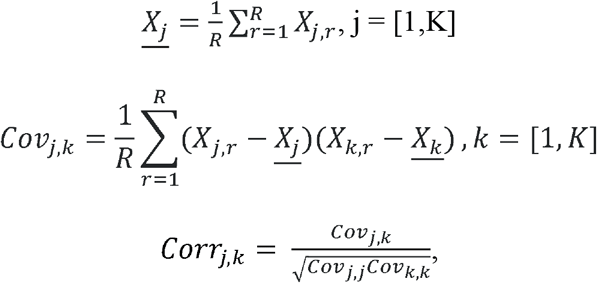

A gene-to-gene co-expression score was considered valid if *Corr_j,k_* > 0.2 and these genes *M* were used in all further analysis. Now, the spatial matrix was subset to create a *R x M* matrix used in the cell typing task and a cell type expression score *c_l,r_* for each gene expression value *Y_m,r_* was calculated:

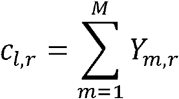

The cell type assignment *c_l, r_* was then scaled between the different cell types present in all the regions:

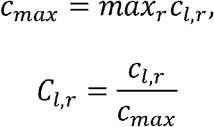

To represent proportions of cell types in each region, we finally scaled the data by the cumulative cell type score calculated for the region such that:

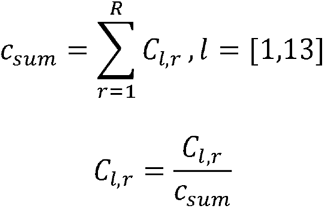

*C_l,r_* represented the approximated contribution of each cell type *I* in each region *r*. The gene signatures *M* were also tested for functional enrichment with Gene Ontology terms with PANTHER^50^. We reported all terms at 5% FDR.

## Data and materials availability

Raw sequencing data is available through an MTA with Vivianne Malmström. All processed data files are available at the Single Cell Portal (https://portals.broadinstitute.org/single_cell/study/SCP460/).

## Code availability

All code has been deposited to https://github.com/klarman-cell-observatory/3dst.

## Acknowledgments

We thank Jennifer Rood for help with manuscript preparation and Ania Hupalowska for illustrations. This research was funded by the Knut and Alice Wallenberg Foundation (S.V., A.C.), the Royal Swedish Academy of Sciences and Swedish Society for Medical Research (S.V.); FOREUM, Foundation for Research in Rheumatology, European Research Council (CoG 2017 - 7722209) and the Konung Gustaf V:s och Drottning Victorias Frimurarestiftelse (A.C.); the Margareta af Ugglas stiftelse (V.M.), the Swedish Rheumatism Association (M.K, V.M.), the Swedish Research Council (V.M., P.L.S), Science for Life Laboratory (P.L.S.) and the Klarman Cell Observatory and HHMI (A.R.). D.S. was supported by the University of Zurich BioEntrepreneur-Fellowship (BIOEF-17-001) and a Swiss National Science Foundation Early Postdoc Mobility fellowship (P2ZHP3_181475); D.S and P.K.S were supported by NCI grant U54-CA225088. S.V. was supported as a Knut and Alice Wallenberg Fellow at the Broad Institute of MIT and Harvard.

## Contributions

S.V. and V.M. designed the study; K.C. performed experiments with supervision from P.L.S.; S.V. designed the analysis and analyzed the data with help from L.L. and B.L.; S.V. implemented the Shiny extension; D.S. designed the histoCAT extension and phenotypic analysis workflows with supervision from P.K.S. and A.R.; M.K. annotated all images; A.I.C. and A.H.H. provided samples for the study; S.V. and A.R. wrote the manuscript with input from all the authors. All the authors read the manuscript and discussed the results.

## Tables

**Supplementary Table 1**. Patient information.

**Supplementary Table 2.** Differentially expressed genes in infiltrate structures (RA1-5).

**Supplementary Table 3.** Differentially expressed genes in all tissue sections (RA1-5).

## Supplementary Figures

**Supplementary Figure 1.**
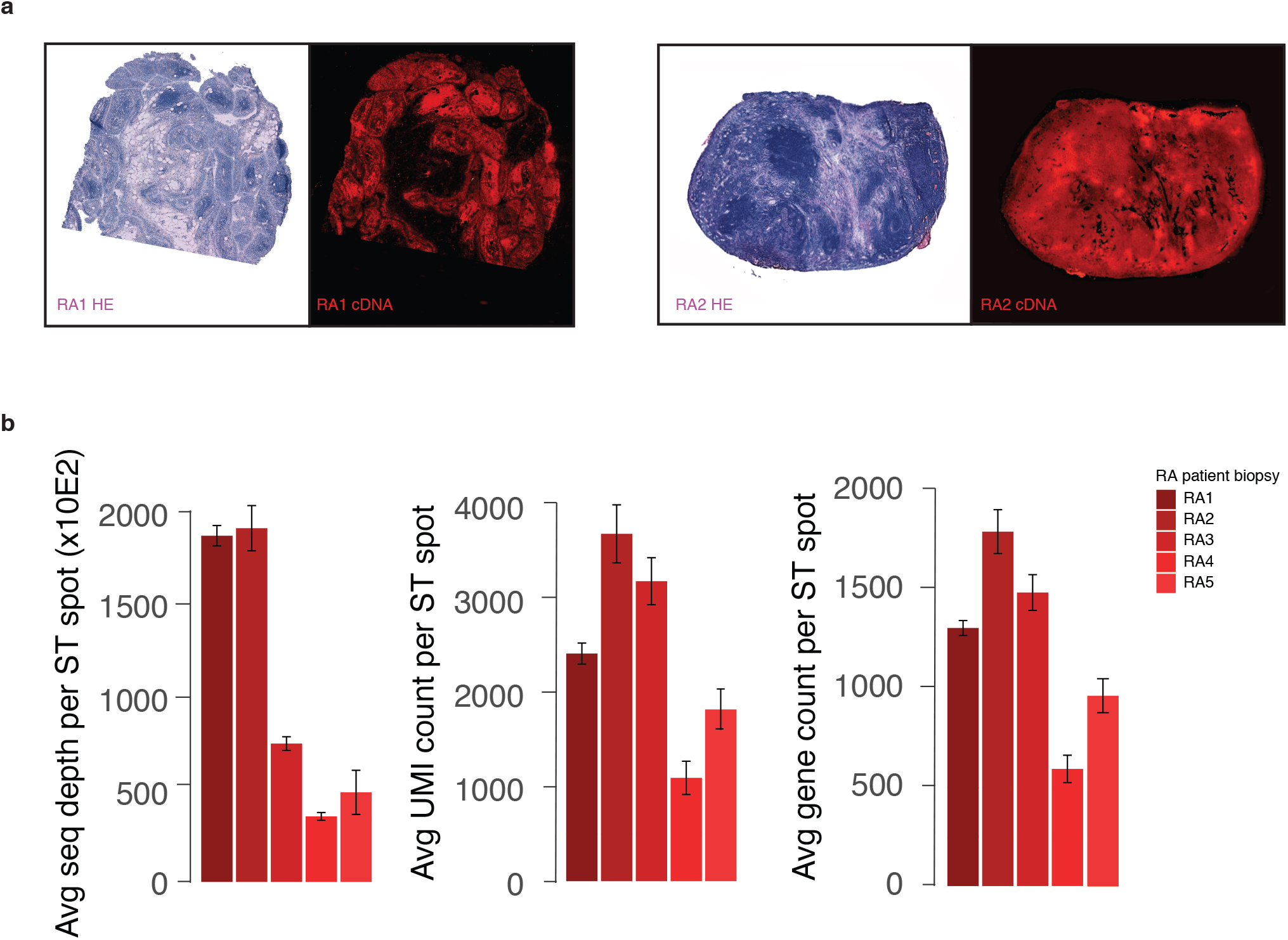
Fluorescent footprint and sequencing library statistics. (**a**) Images of H&E stained tissue sections and corresponding fluorescent cDNA expression footprints marking spatial gene activity for RA1 (knee) and RA2 (hip) patient biopsies. cDNA signal shows optimized tissue handling for both RA sampling sites. (**b**) Sequencing library statistics for all patient biopsies (RA1-5) reporting number of raw sequencing reads, UMIs and unique protein coding gene counts per 100μm spatial feature.

**Supplementary Figure 2.**
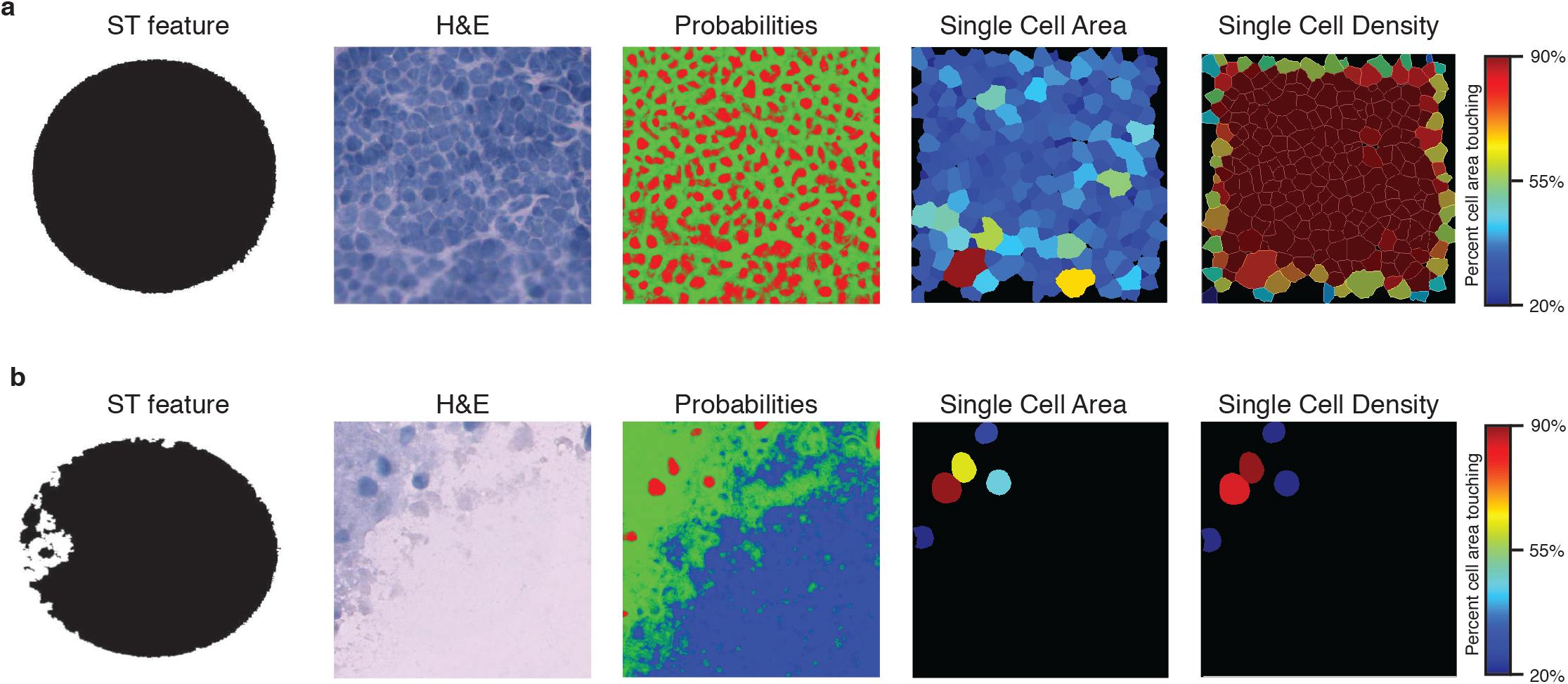
Single cell segmentation and histoCAT analysis workflow. **(a)** 100μm ST features were used to crop the respective H&E images as 100μm x 100μm areas. Ilastik software was used to train a random-forest classifier to create probabilities for three classes in the H&E image (red: nuclei; green: membrane; blue: background). Those probabilities were segmented using the CellProfiler software. Single cell areas (each cell has a discrete color code) and density (color scale represents the percent of each cell’s area touching a membrane of a neighboring cell) were quantified and visualized in the histoCAT software. In these representations, black background color denotes no cells were detected. **(b)** Same as in (a) for a less cell dense area overlapping a 100μm ST feature.

**Supplementary Figure 3.**
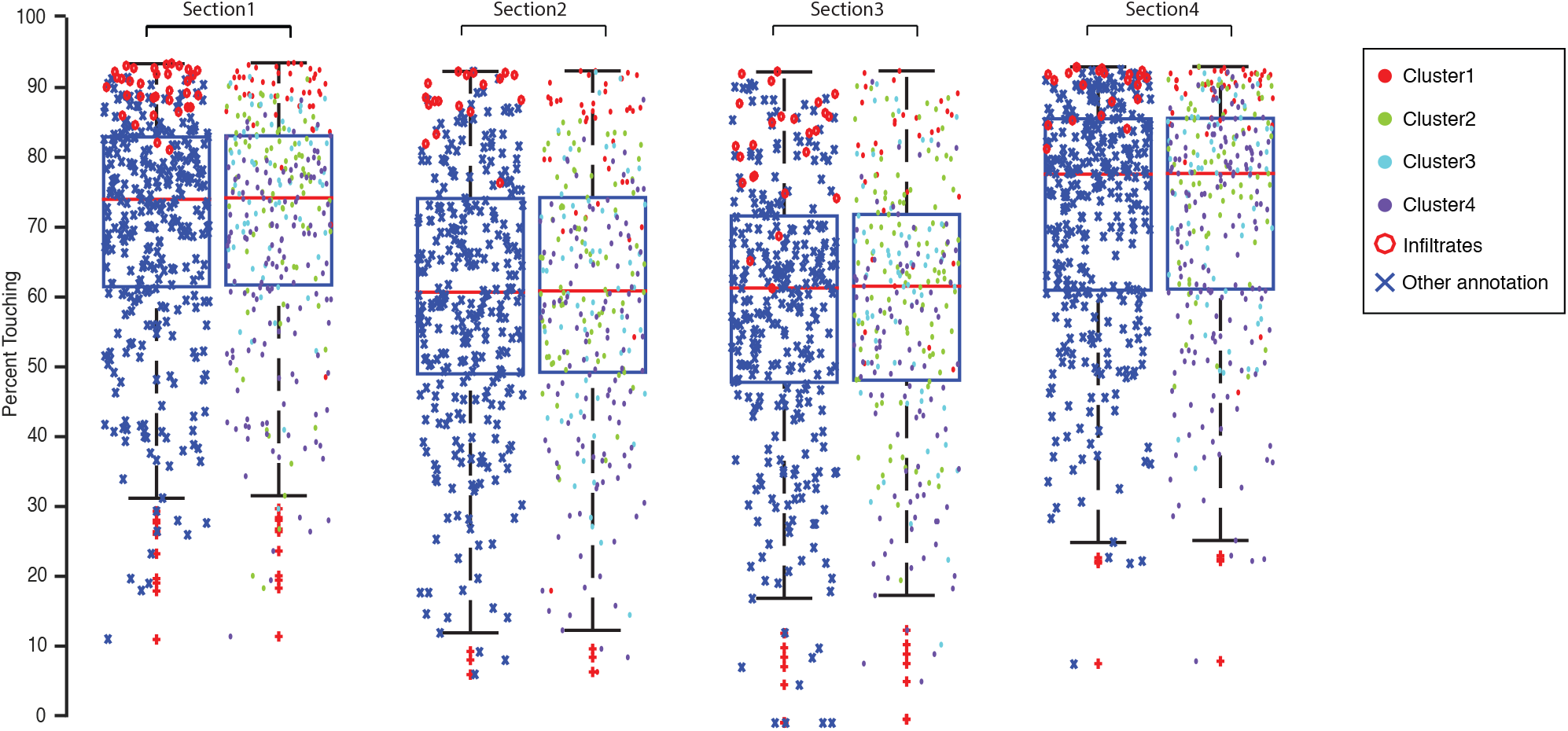
Cellular morphology reproduces manual annotations. On average 80.20% of manually annotated infiltrates (red octagons) were present in ST features with a density higher than 70 in all sections (left boxplot in pair) while at the same density threshold, 91% of all Cluster1 ST features were present in cell dense areas (red circle; right boxplot in pair).

**Supplementary Figure 4.**
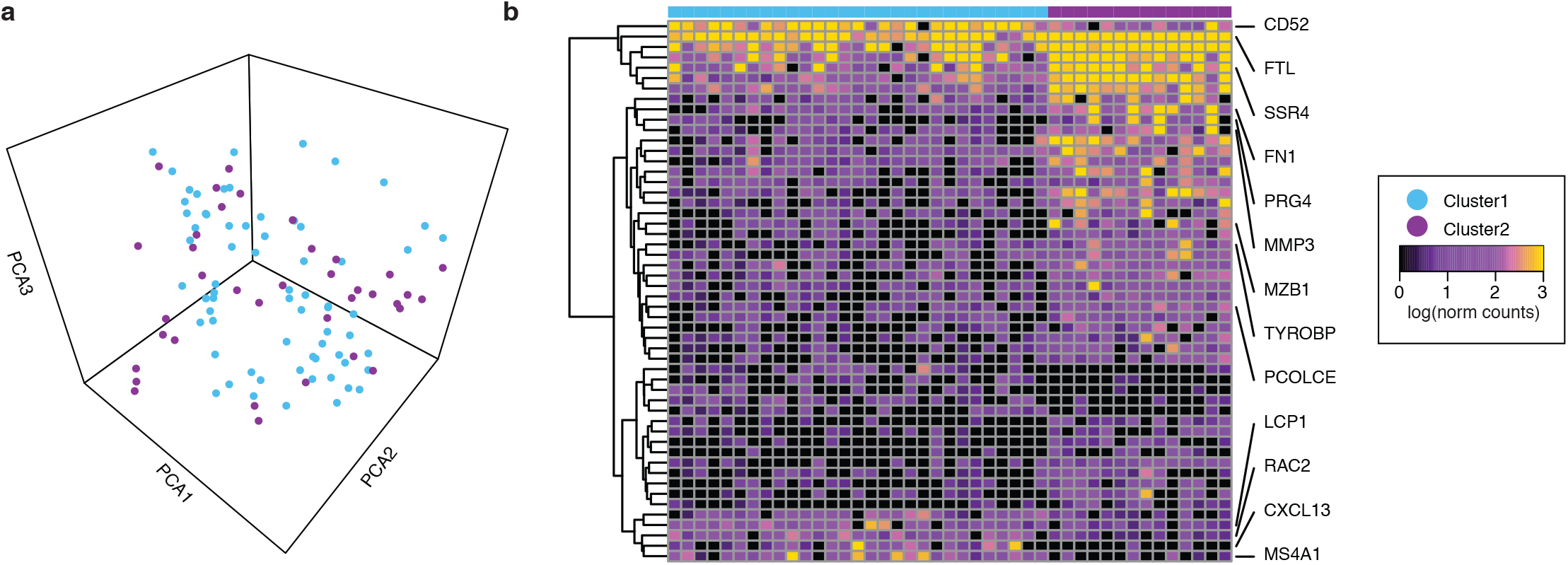
Clustering analysis of RA1 infiltrate regions. (**a**) PCA plot of each individual spatially resolved infiltrate feature present in any of the RA1 tissue sections. Samples have been color-coded based on hierarchical clusters (cyan; purple). (**b**) Heatmap of differentially expressed genes (color scale, rows) between the two clusters (color code, columns) as determined in (a). Color code is shared between the panels.

**Supplementary Figure 5.**
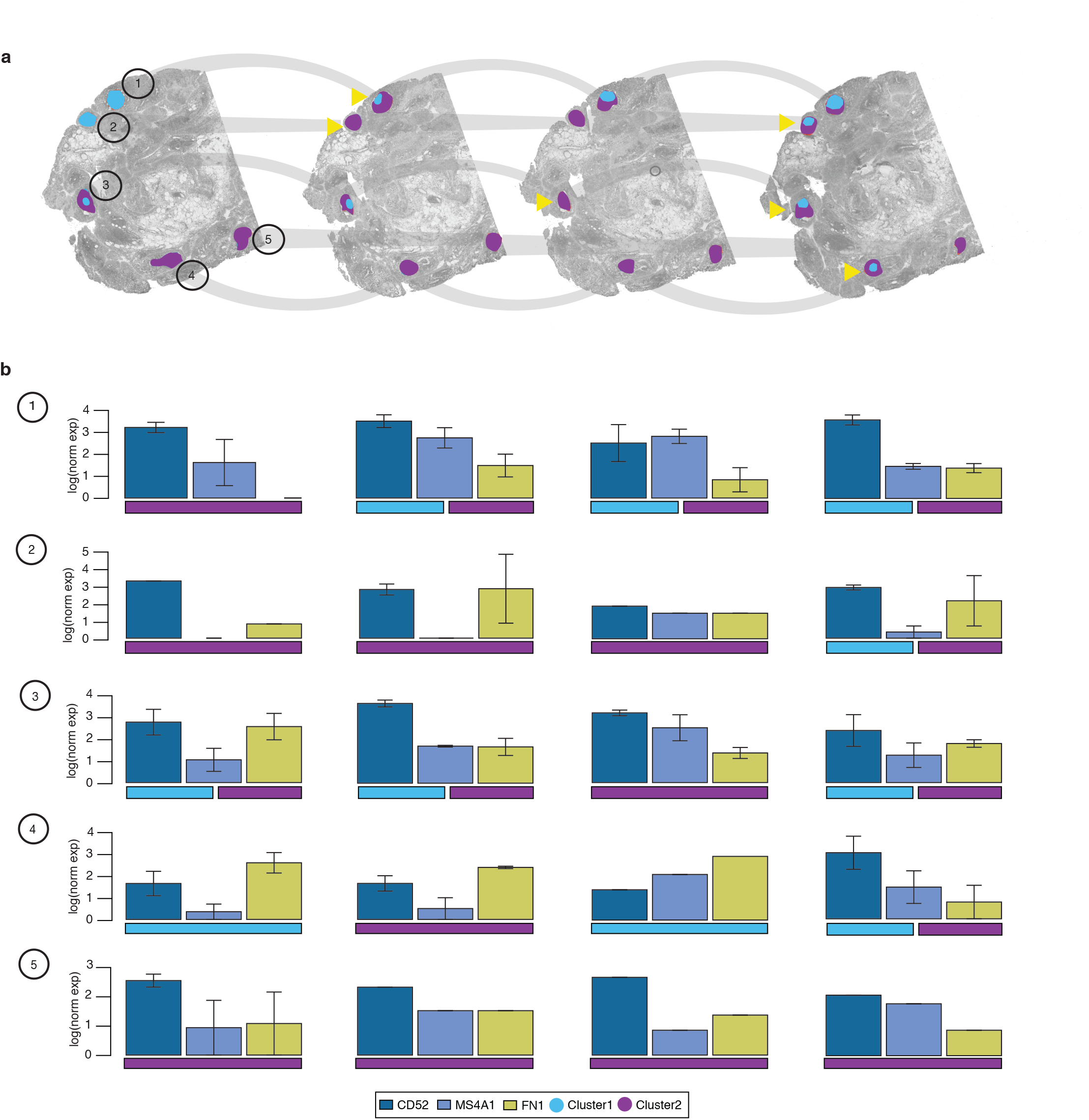
Spatial averages of annotated infiltrates in four RA1 biopsy sections. (**a**) Volumetric morphological view with overlaid color-coded infiltrate regions present in all four sections as determined by clustering (color code). Yellow arrow marks an event where the infiltrate region changed its cluster assignment. (**b**) Barplots showing average expression of three cluster driving genes; CD52; MS4A1 and FN1. Infiltrate regions are number coded (1-5) and the numbering is shared between the panels. Error bars represent s.e.m. where more than one spatial data point was present.

**Supplementary Figure 6.**
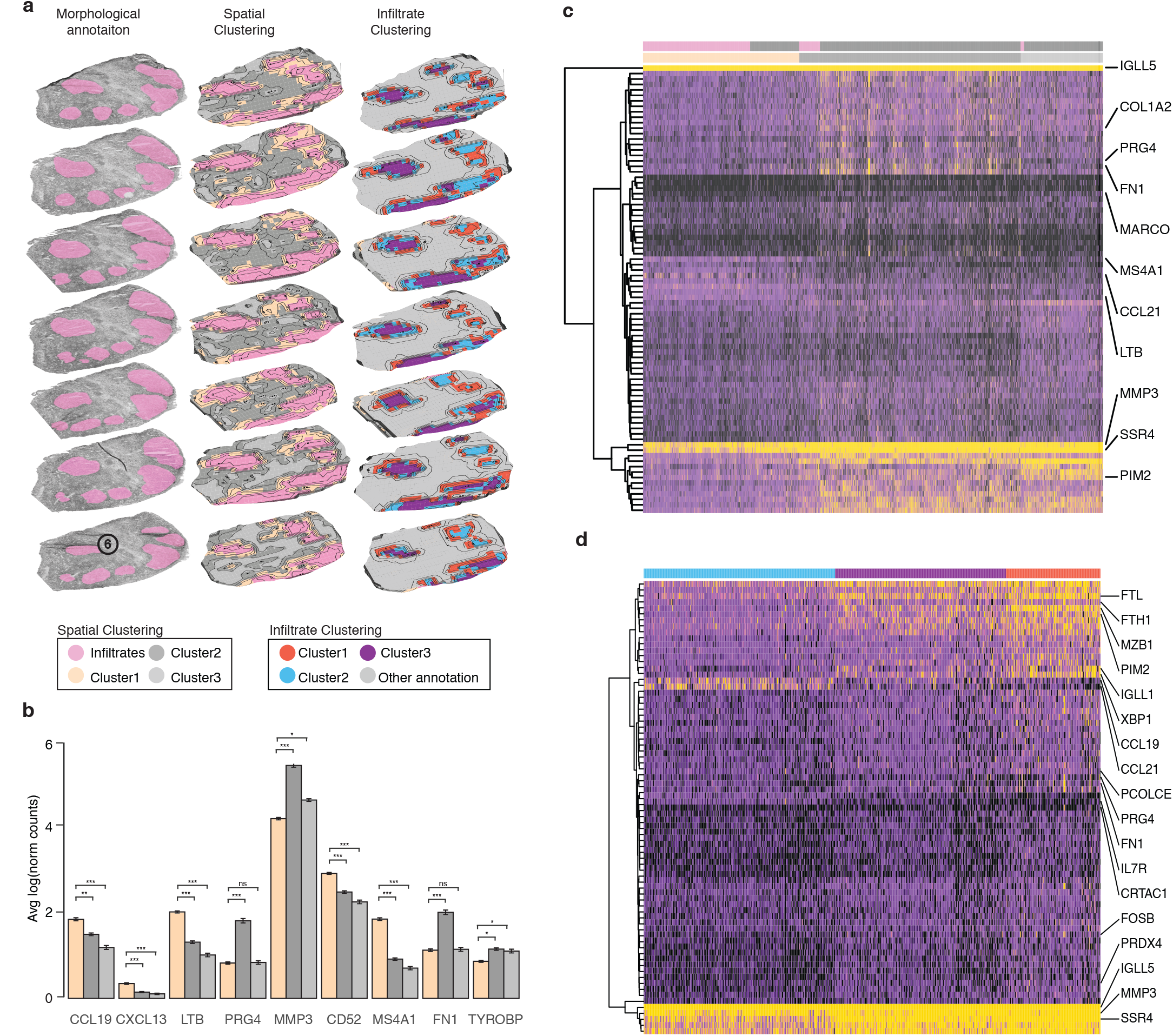
Spatial gene expression in the RA2 patient biopsy. (**a**) Spatial clustering (color code) as compared to morphological annotation and infiltrate clustering (color code). Pink (color code) marks infiltrate regions that we found as overlapping between the morphological annotation and spatial clusters (Cluster 1). Color codes are shared between the panels. (**b**) Average expression of some spatially variable genes in the clusters. Wilcoxon’s rank sum test for PRG4, MMP3, FN1 and TYROBP denotes difference in Cluster 1 lesser than in another cluster while for the rest of the genes, the same denotation describes differences greater in Cluster 1 than the rest. Statistical significance markings (Wilcoxon’s rank sum test) are displayed; p>0.5 (ns), 0.005<p≤0.05(*), 0.0005<p≤0.005(**), p≤0.0005(***). **(c)** Heatmap of gene expression (color scale) where each column represents one spatial feature and each row a gene. Spatial features (columns) have been color-coded in the top panel into two annotation categories (pink; annotated infiltrates and dark grey; rest other annotation). In the bottom annotation panel, spatial features were color coded based on their spatial cluster identities as determined in (a). Example genes (rows) have been highlighted in the image. (**d**) Heatmap of differentially expressed genes (color scale, rows) between the three infiltrate clusters (color code, columns) as determined by infiltrate clustering in (a). Example genes (rows) have been highlighted in the image.

**Supplementary Figure 7.**
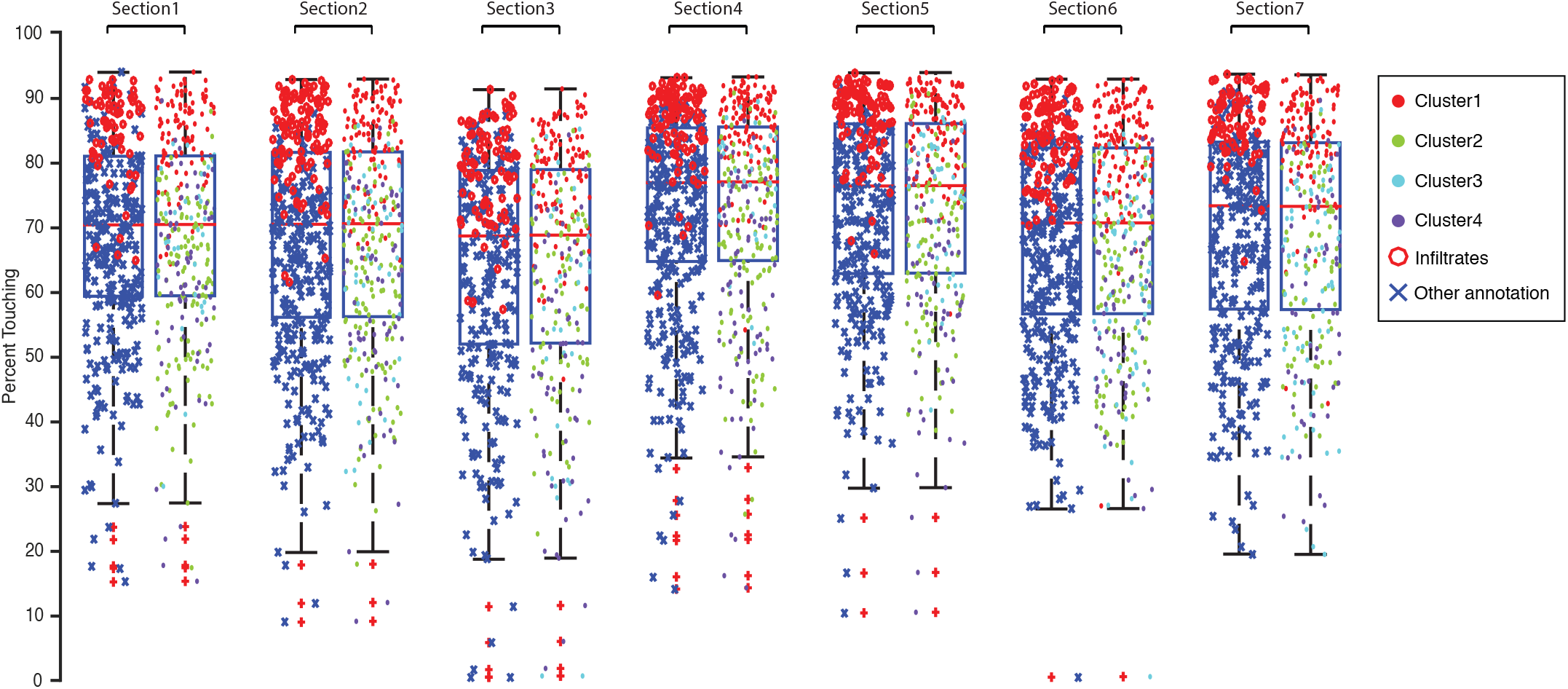
Cellular morphology improves manual annotation in RA2 patient sections. On average 90% of manually annotated infiltrates (red octagons) were present in ST features with a density higher than 70% in all sections (left boxplot in pair) while at the same density threshold, 84% of all Cluster1 ST features were present in cell dense areas (red circle; right boxplot in pair).

**Supplementary Figure 8.**
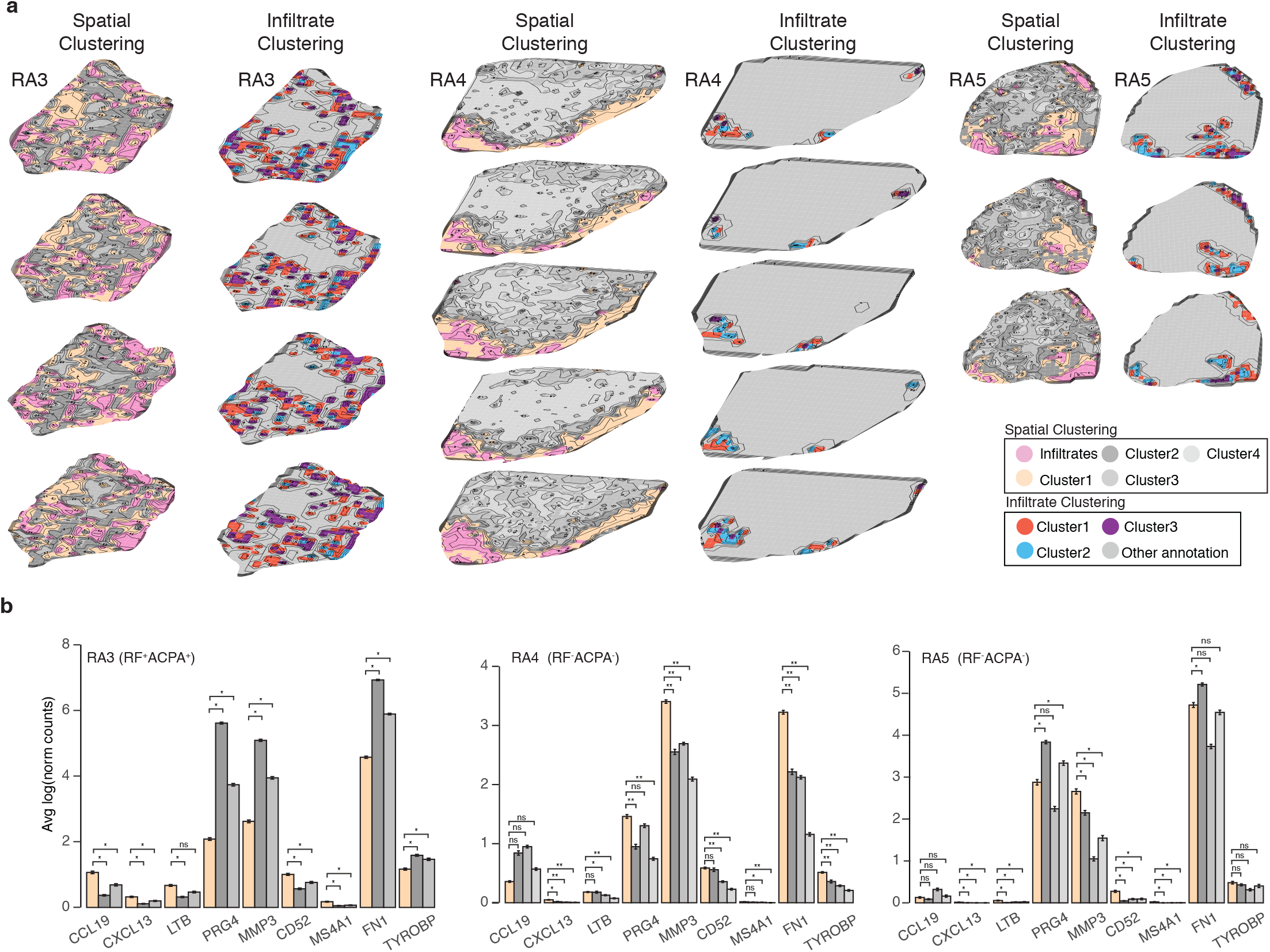
Spatial and infiltrate clustering for RA3-5 patient biopsies. (**a**) Spatial clustering (color code) as compared to clustering of only infiltrate regions (color code). Pink (color code) marks infiltrate regions that we found as overlapping between the morphological annotation and spatial clusters (Cluster1). (**b**) Average expression of some spatially variable genes in the clusters. For RA3, Wilcoxon’s rank sum test for PRG4, MMP3, FN1 and TYROBP denotes difference in Cluster 1 lesser than in another cluster while for the rest of the genes, the same denotation describes differences greater in Cluster 1 than the rest. For RA4, all differences are described as greater while for RA5; PRG4, FN1 and TYROBP are only genes that denote lesser significant expression in Cluster 1. Statistical significance markings (Wilcoxon’s rank sum test) are displayed; p>0.5 (ns), 0.005<p≤0.05(*), 0.0005<p≤0.005(**), p≤0.0005(***).

**Supplementary Figure 9.**
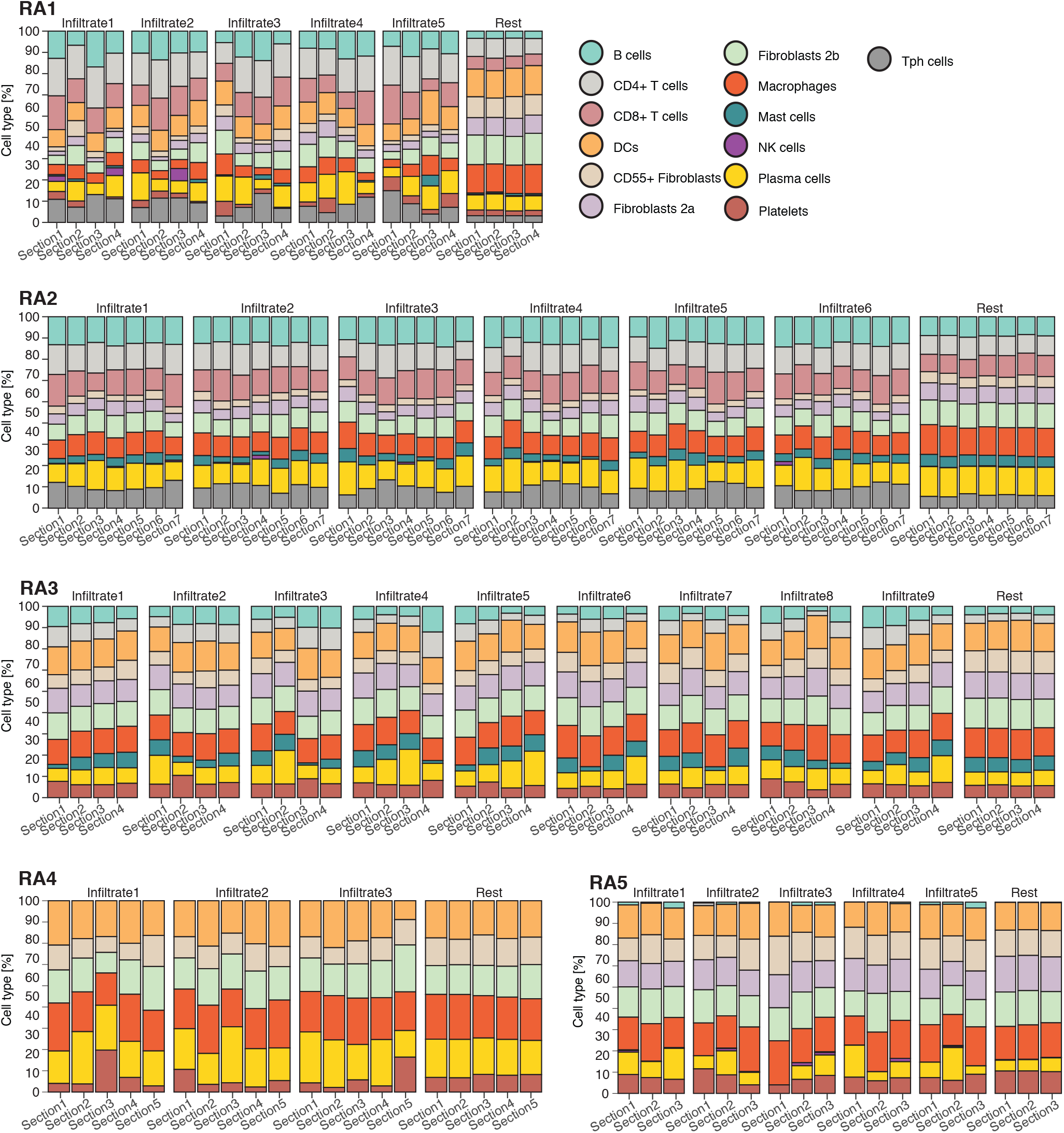
Single cell distributions in the tissue volume. Cell type percentage of 13 tested cell types (color code) shown in each section (column) and divided in groups of TLO infiltrates or the rest of the tissue (“Rest”).

**Supplementary Figure 10.**
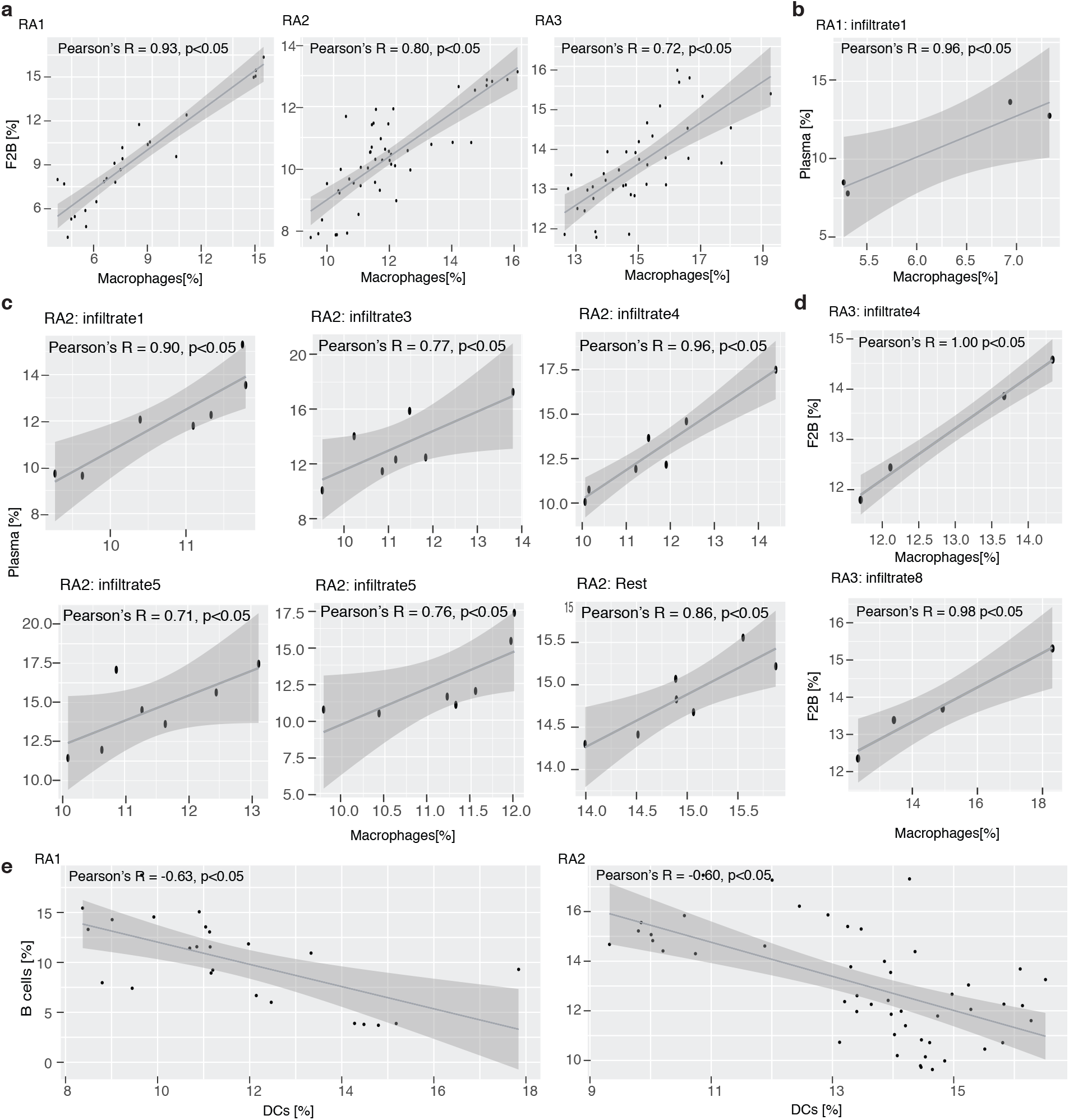
Correlations between different spatial cell type abundances in the rheumatoid arthritis synovium. (**a**) Correlation plots between macrophage and F2B abundances in the whole tissue volume in RA1-3. (**b-c**) Correlation plots between macrophage and plasma cell abundances in infiltrates or surrounding regions (“rest”) in RA1 and RA2. (**d**) Correlation plots between macrophage and F2B abundances in two infiltrate regions in RA3. (**e**) Correlation plots between dendritic cell (DCs) and F2B abundances in the whole tissue volume in RA1 and RA2. Reported are Pearson’s correlation coefficients (R) and empirical p values for each comparison.

**Supplementary Figure 11.**
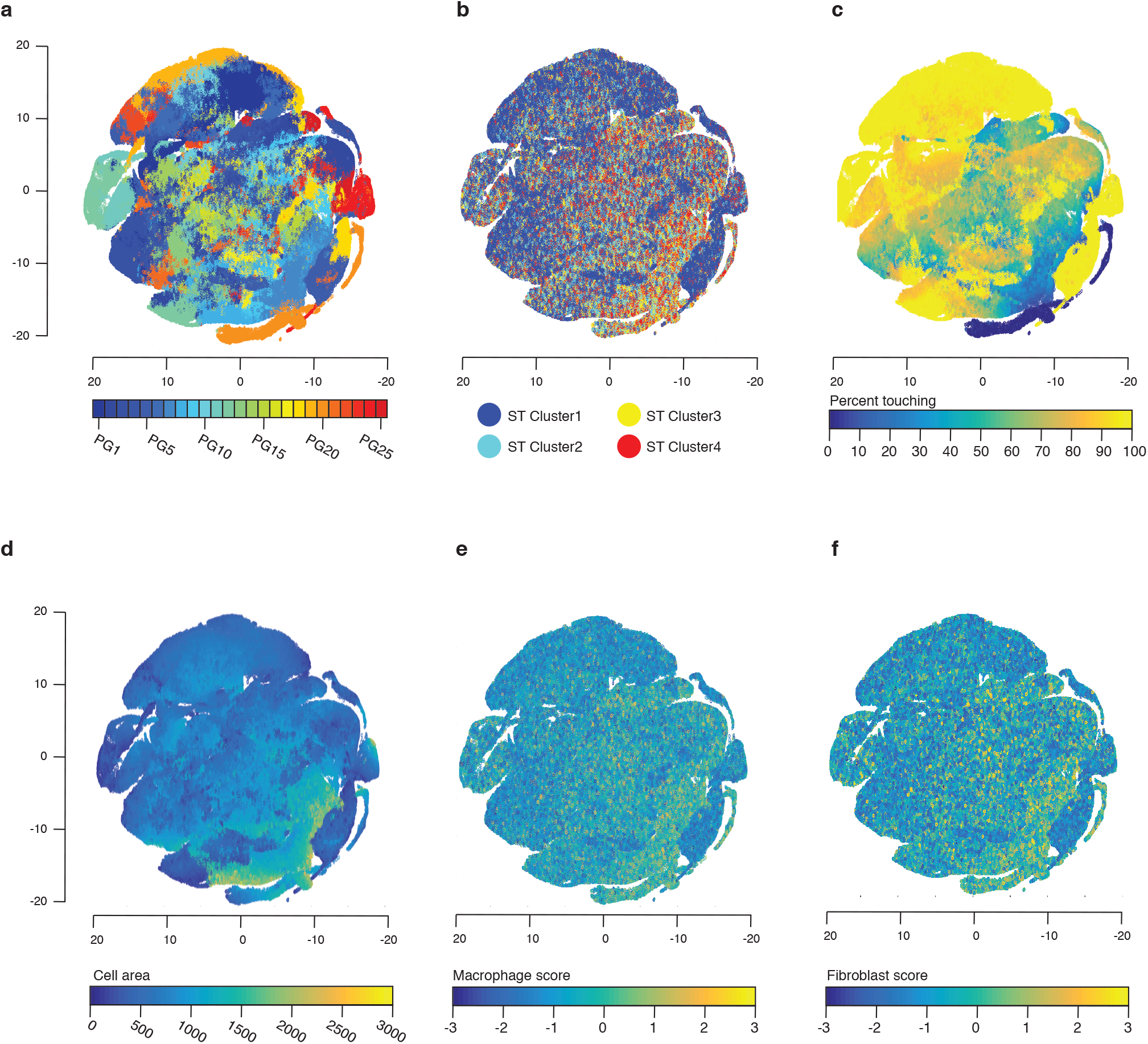
Dimensionality reduced RA2 topological features correlate with cell type expression. **(a)** 25 phenotypic groups (PG) clustering visualized in tSNE projection. **(b)** ST cluster color codes overlaid on top of (a). **(c)** Cell density as percent cells touching another cell overlaid on top of (a). **(d)** Cell area overlaid on top of (a). **(e)** Macrophage cell type score overlaid on top of (a). **(f)** Fibroblast (F2B) cell type score overlaid on top of (a).

